# Tetrabromobisphenol A (TBBPA) inhibits *Bacillus subtilis* as a Membrane Active Antibacterial Agent

**DOI:** 10.64898/2026.02.10.705026

**Authors:** Fang Ji, Zhunjie Li, Yusheng Wang, Jurian Wijnheijmer, Leendert W Hamoen

## Abstract

Antimicrobial resistance (AMR) is a pressing global public health crisis, necessitating novel antimicrobial agents and mechanistic insights. Tetrabromobisphenol A (TBBPA), a widely used brominated flame retardant, exhibits potent activity against Gram-positive bacteria including methicillin-resistant *Staphylococcus aureus* (MRSA) without inducing resistance, yet its mode of action remains unclear. Using *Bacillus subtilis* as a model, we investigated TBBPA’s antibacterial mechanism via extensive bacterial cytological profiling, fluorescence imaging, and mutant validation. We found that TBBPA causes membrane depolarization and disruption, based on Thioflavin T release, MinD mislocalization, FM5-95 fluorescence aggregation, and propidium iodide penetration. In fact, TBBPA can destabilizes giant unilamellar lipid vesicles *in vitro*. The induced membrane damage triggers several downstream effects, including MreB immobilization, which impairs cell wall synthesis, inhibition of DNA replication and translation, and increased autolysin activation leading to cell lysis. In conclusion, this study suggests that TBBPA directly targets the cell membrane, causing disruption of multiple essential processes, and leads to activation of autolysins, resulting in lysis. These findings highlight TBBPA’s prospective utility as an anti-Gram-positive agent; nevertheless, concerns over its potential side effects necessitate further investigations into its safety profile prior to clinical translation.

## INTRODUCTION

Antimicrobial resistance (AMR) is a significant global concern, posing one of the top public health threats. In 2019, it was estimated that there were approximately 4.95 million deaths associated with bacterial AMR, including 1.27 million directly attributable to this issue [1]. To address the pressing antimicrobial resistance crisis, the discovery of novel antimicrobial compounds, and understanding their mechanisms, is urgently necessary.

In a previous study, we have demonstrated the antimicrobial efficacy of the bromated flame retardant Tetrabromobisphenol A (TBBPA) (Fig. 1A), against Gram-positive bacteria, including Methicillin-resistant *Staphylococcus aureus* (MRSA), without detecting resistance [2]. Bromated flame retardants have been widely used to decrease the flammability of various commercial products, and TBBPA stands out as one of the most commonly used flame retardants worldwide in a variety of consumer products, with approximately 90 % use in manufacturing resins [3-5]. Owing to the high volume of production and usage, the environmental toxicity of TBBPA has been investigated in depth. At concentrations in the μM range TBBPA shows acute toxicity to aquatic organisms [4, 6]. Numerous *in vitro* and *in vivo* studies utilizing cell and animal models have illustrated that TBBPA is capable of inducing pleiotropic effects in both cells and animals, with potential toxicological impacts on multiple organs and systems, including the liver, kidneys, nervous system, heart, endocrine and reproductive system [7-11]. This compound is classified as a group 2A carcinogen (likely carcinogenic to humans) by the International Agency for Research on Cancer [12]. However, data on human toxicity is scarce [13, 14]. According to a European Union Risk Assessment Report, no health effects of concern have been identified for TBBPA in humans [15]. Low levels of human exposure to TBBPA coupled with its rapid metabolic clearance in the human body might suggest that TBBPA could be a relatively safe chemical for the general population[16].

**Fig. 1.**
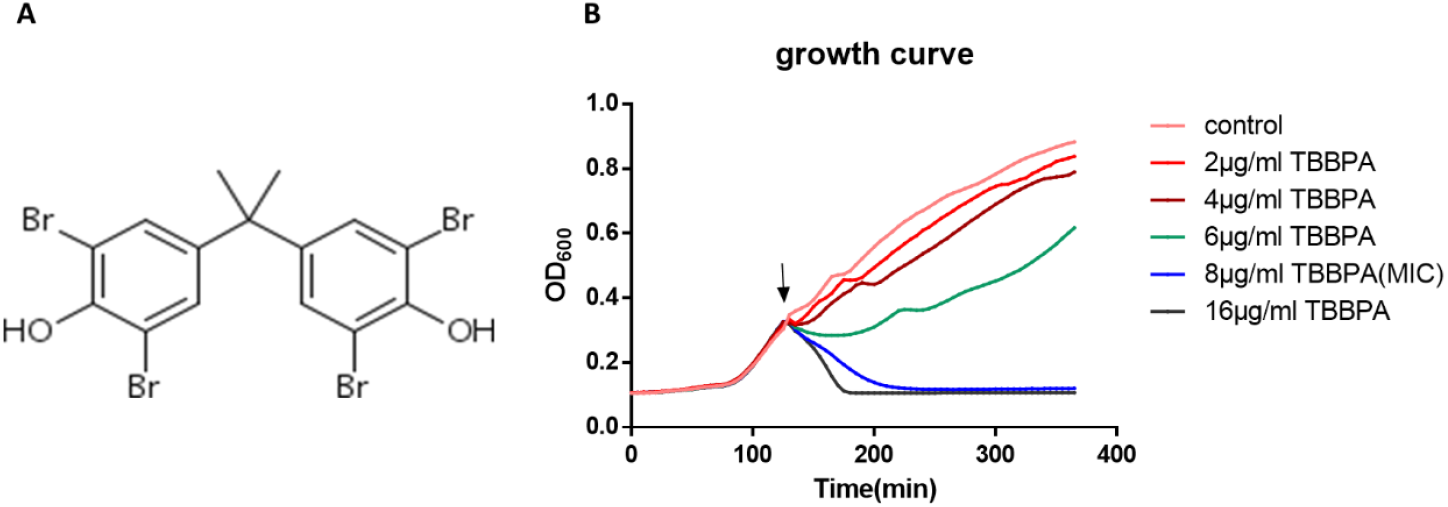
Impact of TBBPA on the growth of *B. subtilis*. (A) Molecular structure of TBBPA. (B) Inhibitory effect of TBBPA on exponentially growing *B. subtilis*. The arrow marks the time point of TBBPA addition.

The inability to isolate resistant mutants positions TBBPA as an appealing candidate for clinical application as an antibiotic, However, it complicates the determination of its mechanism of action by genetic screening. TBBPA specifically targets Gram-positive bacteria and we have shown that it induces lysis of the Gram-positive model system *Bacillus subtilis*, by a yet unknown mechanism [2]. A transcriptome analysis revealed that the *dlt* gene cluster and two autolysin genes, *yocH* and *cwlO*, were upregulated. The *dlt* operon is responsible for the addition of D-alanyl to teichoic acids, which are uniquely present in the cell walls of Gram-positive bacteria. It has been speculated that increased D-alanylation influences the number of anionic sites on teichoic acids, thereby reducing autolysin binding and less autolysis [17]. TBBPA exposure also results in the upregulation of three peptidoglycan synthesis-related genes, *murB, ponA*, and *pbpA*, which encode UDP-N-acetylenolpyruvoylglucosamine reductase, penicillin-binding protein (PBP) 1 and penicillin-binding protein 2A, respectively [18-20]. This genetic information suggested that TBBPA targets a crucial cell wall synthesis step, leading to cell wall degradation and cell lysis, possibly stimulated by the increased expression of autolysin genes. To gain a better understanding of the exact mode of action of TBBPA, we performed extensive bacterial cytological profiling on TBBPA exposed *B. subtilis* cells, a technique that has been successfully used to elucidate the killing mechanism of antibacterial compounds [21, 22]. This study revealed that TBBPA targets directly the cell membrane, leading to membrane depolarization, thereby activating autolysin activity and cell lysis. The consequences of this mode of action are further discussed.

## MATERIAL AND MEHODS

### Antimicrobial agents and chemicals

TBBPA was purchased from Sigma. Daptomycin was purchased from Novartis. All other antibiotics were purchased from Sigma Aldrich in the highest possible purity. TBBPA, gramicidin, gramicidin S, CCCP and valinomycin were dissolved in sterile DMSO. Vancomycin, ciprofloxacin, and nisin were dissolved in sterile water. Chloramphenicol, erythromycin, and rifampicin were dissolved in ethanol.

### Bacterial strains and growth conditions

*B. subtilis* strains used in this study are listed in Table S1. Unless otherwise noted, all strains were grown in Luria Bertani (LB) broth at 37°C or 30°C (for microscopy) under steady agitation in the presence of inducer where appropriate (see Table S1). All experiments were performed in early exponential growth phase (OD_600_ = 0.3–0.35). Unless otherwise stated, experiments were carried out in biological triplicates.

### Minimal inhibitory concentration (MIC) assay and growth curve

The MICs of TBBPA was determined by the standard micro-dilution method recommended by the Clinical and Laboratory Standards Institute [23]. Growth experiments were performed with a Biotek Synergy MX plate reader in 96-well format under continuous shaking in a final volume of 150 μl per well. *B. subtilis 168* was grown in LB at 37°C until an OD_600_ of 0.3 and subsequently treated with 2, 4, 6, 8 and 16 μg/ml TBBPA (0.25x, 0.5x, 0.75x, 1x, and 2x MIC), or DMSO as control.

### Localization of fluorescent fusion proteins

Strains expressing fluorescent fusion proteins were grown at 30°C in the presence of appropriate inducer concentrations (S1 Table) until an OD_600_ of 0.3. Cells were subsequently treated with 8 μg/ml (1x MIC) TBBPA or 1% DMSO (negative control). Unless otherwise stated, samples for microscopy were withdrawn after 15min of treatment. Cells were immobilized on a thin film of 1% agarose and immediately observed using an Olympus BX 50 microscope equipped with a Photometrics CoolSNAP fx digital camera. Images were analyzed with Image J.

### Thioflavin T (ThT) assay

The membrane potential was determined with the potentiometric fluorescent probe Thioflavin T (ThT) using a Biotek Synergy MX plate reader. In short, *B. subtilis* 168 overnight cultures were washed and diluted to OD_600_ 0.08 with buffer (10 mM K_3_PO_4_, 5 mM MgSO4, and 250mM sucrose (pH 7.0)), the baseline was recorded (450 nm excitation, 482 nm emission). 10μM Tht was then added and the fluorescence signal was recorded for 5 min. Compounds were added (8μg/ml (1x MIC) TBBPA, 0.5μM CCCP (1x MIC), or 1% DMSO) with CCCP as positive control and DMSO as negative control, and samples were measured for another 20 min (37°C, shaking).

### Propidium iodide (PI) pore staining

Exponentially growing (37°C) *B. subtilis 168* cultures were treated with 16 μg/ml (2x MIC) TBBPA, 0.5% SDS (positive control), or 1% DMSO (negative control). Samples were withdrawn after 15min of treatment and stained with 10 μg/ml PI for 5 min in the dark (37°C, shaking). Cells were washed twice with pre-warmed phosphate buffered saline (PBS) and resuspended in the same buffer. PI fluorescence was measured using a Biotek Synergy MX plate reader (535 nm excitation, 617 nm emission).

### Membrane staining

All membrane-specific experiments were performed at 30°C. Unless otherwise stated in figure legends, exponentially growing cells were treated with 8μg/ml (1x MIC) TBBPA for 10 min. Membranes were stained with 2μg/ml FM5-95 (Molecular Probes) for 10 min immediately prior to microscopy. Nucleoids were stained with 1μg/ml DAPI (Thermo Scientific) for 2 min immediately prior to microscopy.

### Giant unilamellar vesicles (GUVs) assay

GUVs were prepared as below by modifying the gel-assisted method described previously [24]. A 5% (w/w) solution of polyvinyl alcohol (PVA, MW 145,000, VWR International) was prepared by mixing PVA with water and heating it in a microwave. The PVA gel was allowed to cool to room temperature before use. Approximately 200 μl of the PVA gel was applied to a glass slide, where it was spread thinly and evenly, and the PVA gel thin layer was then dried at 55°C for about 30 min. *E. coli* polar lipid extract (product and supplier information) was dissolved in chloroform to 1mg/ml. About 40-60 μl of the lipid dissolved in chloroform was spread onto the PVA film and then dried using argon stream to remove chloroform and form a dry lipid thin layer. To generate GUVs, 40-100 μl of 25 mM Tris-HCl buffer (pH 7.5), supplemented with 280 mM sucrose and 150 mM K_2_SO_4_, was added to cover the lipid thin layer. The formation of GUVs happened after 15-30 min incubation at room temperature. The liquid layer containing GUVs was transferred to Eppendorf tubes using a blunt-tipped 200 μl pipette and left at room temperature for 10-15 min. The GUV suspension was then slowly diluted (1:20) in a 25 mM Tris-HCl buffer (pH 7.5), supplemented with 280 mM glucose and 150 mM Na_2_SO_4_ to allow sedimentation, followed by staining with 1 μg/ml Nile Red. 47.5 μl of the diluted GUVs suspension was transferred to homemade wells. To create the homemade wells, the rings from 1 ml tips were cut and attached to a piece of cover glass with glue, which was allowed to dry overnight at room temperature. Before microscopy, an additional microscope slide-shaped iron frame was used to fix the cover glass with wells so that the entire setup could be compatible to the microscope. After loading the samples, the plate was left in the dark at room temperature for 15 min until all GUVs had settled at the bottom of the plates. Thereafter 2.5 μl of chemical solution was added to a well, and the GUVs were observed and imaged using structured illumination microscopy (SIM). The GUVs were treated with either 80 μg/mL TBBPA, 2.5% DMSO (negative control), or 0.2% SDS (positive control). 3D SIM imaging was performed using a Nikon Eclipse Ti N-SIM E microscope setup equipped with a CFI SR Apochromat TIRF 100x oil objective (NA 1.49), a LU-N3-SIM laser unit, an Orca-Flash 4.0 sCMOS camera (Hamamatsu Photonics K.K.), and NIS Elements AR software.

### Transmission electron microscopy

*B. subtilis 168* was grown in LB until an OD_600_ of 0.6-0.8 and subsequently treated with 16 μg/ml (2x MIC) TBBPA or equal amount of DMSO as control. After 30 min of treatment. the cultures were centrifuged and prepared for transmission electron microscopy (TEM). TEM was performed following the method described by Zhu *et al* [25].

## RESULTS

### Bacterial cytological profiling

Bacterial cytological profiling is a phenotypic analysis technique that looks at individual cells utilizing fluorescence microscopy and an array of different fluorescent markers [21, 22]. To determine the proper TBBPA concentrations and incubation times to be able to observe relevant cellular changes with bacterial cytological profiling, we tested different concentrations of TBBPA around the minimum inhibitory concentration (MIC) of 8 μg/ml by following the optical density (OD) (Fig. 1B). At MIC concentrations and above, the OD values decreased quickly, indicating rapid cell lysis. Slightly lower concentrations of 6 μ g/ml caused some growth retardation but did not result in a clear lysis of the culture, and 4 μg/ml and lower did not show a clear effect on growth. Based on this information we exposed cells to 8 μg/ml TBBPA for only 15 min, approximately one-third of the average cell cycle [26] for cells grown at 30 °C.

First, we examined the effect of TBBPA on DNA replication, transcription, translation, and possible DNA damage using reporter strains containing GFP-labeled subunits of DNA polymerase (DnaN), RNA polymerase (RpoC), ribosomes (RpsB) and a GFP fusion with the DNA repair protein RecA (Fig. 2). DNA polymerase forms clear foci in the cell when active [27], this clustering was annihilated following TBBPA exposure. The effect on RNA polymerase localization is less apparent, but the distribution of ribosomes changed as well and became more equally dispersed throughout the cell, which is observed when translation is blocked [28]. Upon DNA damage, RecA forms large nucleoprotein filaments [29], however, such features were not evident in our study. These data indicate that the presence of TBBPA blocks DNA replication but not due to DNA damage, and that protein translation is also likely affected.

**Fig. 2.**
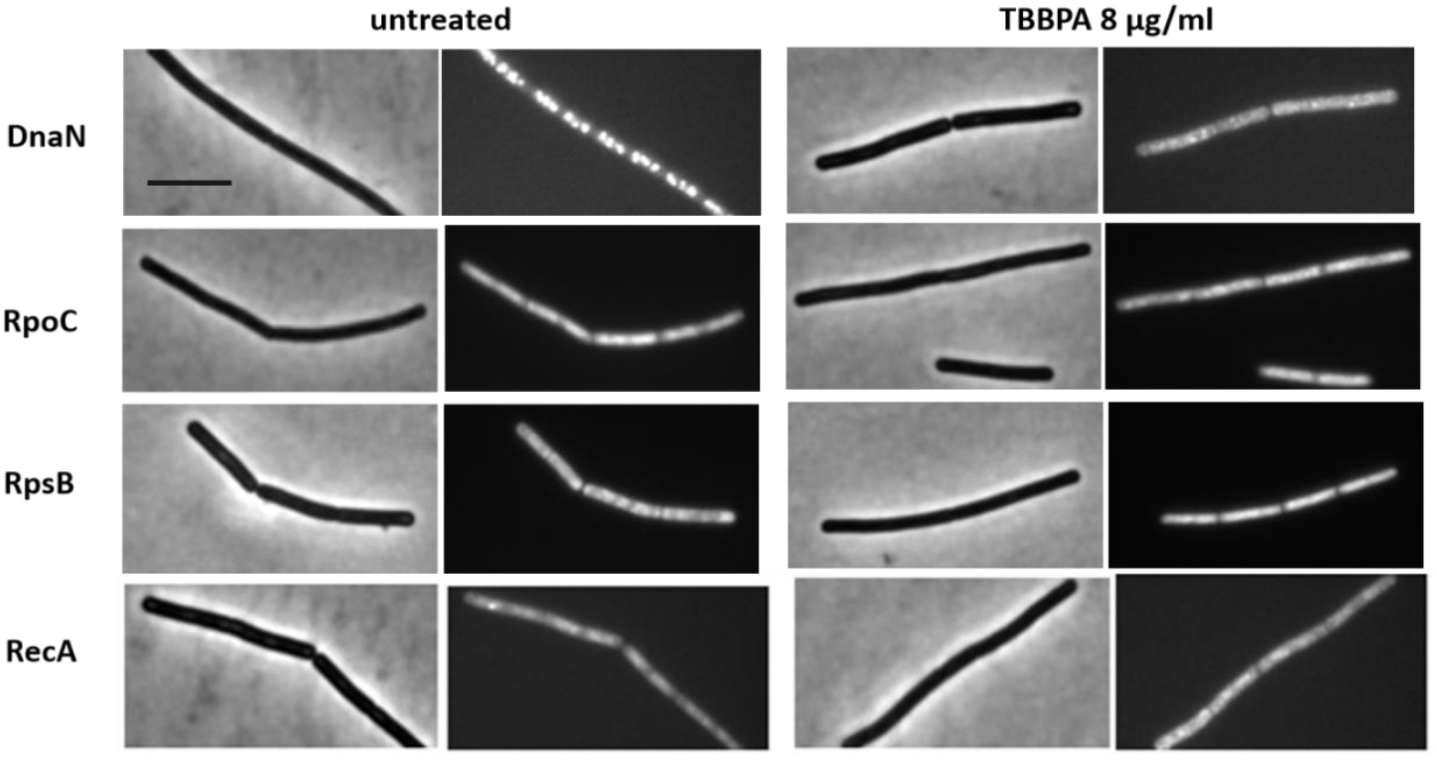
Localization of intracellular reporter proteins. *B. subtilis* strains (see S1 Table for detailed information) HM771 (*dnaN*-gfp, reporter for impaired replication), 1048 (*rpoC*-gfp, reporter for impaired RNA synthesis), 1049 (*rpsB*-gfp, reporter for impaired protein synthesis) and UG-10 (*recA*-gfp, reporter for DNA damage), were grown until an OD_600_ of 0.3 and subsequently treated with 8 μg/ml TBBPA. Scale bar 2μm.

### Effect of TBBPA on MreB mobility

The effect of TBBPA on DNA replication and translation cannot explain the rapid cell lysis. Therefore, we investigated whether TBBPA also affects cell wall synthesis by looking at the mobility of the actin homolog MreB. MreB polymerizes into filaments along the cell membrane [30], and plays a crucial role in organizing lateral cell wall synthesis in many rod-shaped bacteria. MreB and the elongation machinery move circumferentially around the cell in a process driven by cell wall synthesis [31-33]. Exposure to cell wall synthesis inhibitors blocks this movement [8], which can be monitored by taking fluorescent images of GFP-tagged MreB with short time intervals [34]. In this experiment, we used a slightly lower concentration of TBBPA, 5 µg/mL, to reduce the chance of cell lysis. As shown in Fig. 3, GFP-MreB foci move over a time interval of 30 sec. Blockage of peptidoglycan synthesis with either nisin or vancomycin, which target the peptidoglycan precursor lipid II, and the D-Ala-D-Ala terminus of peptidoglycan, respectively [35, 36], stops this movement, as has been shown before [34]. Interestingly, sub-MIC concentrations of TBBPA gave the same effect (Fig. 3).

**Fig. 3.**
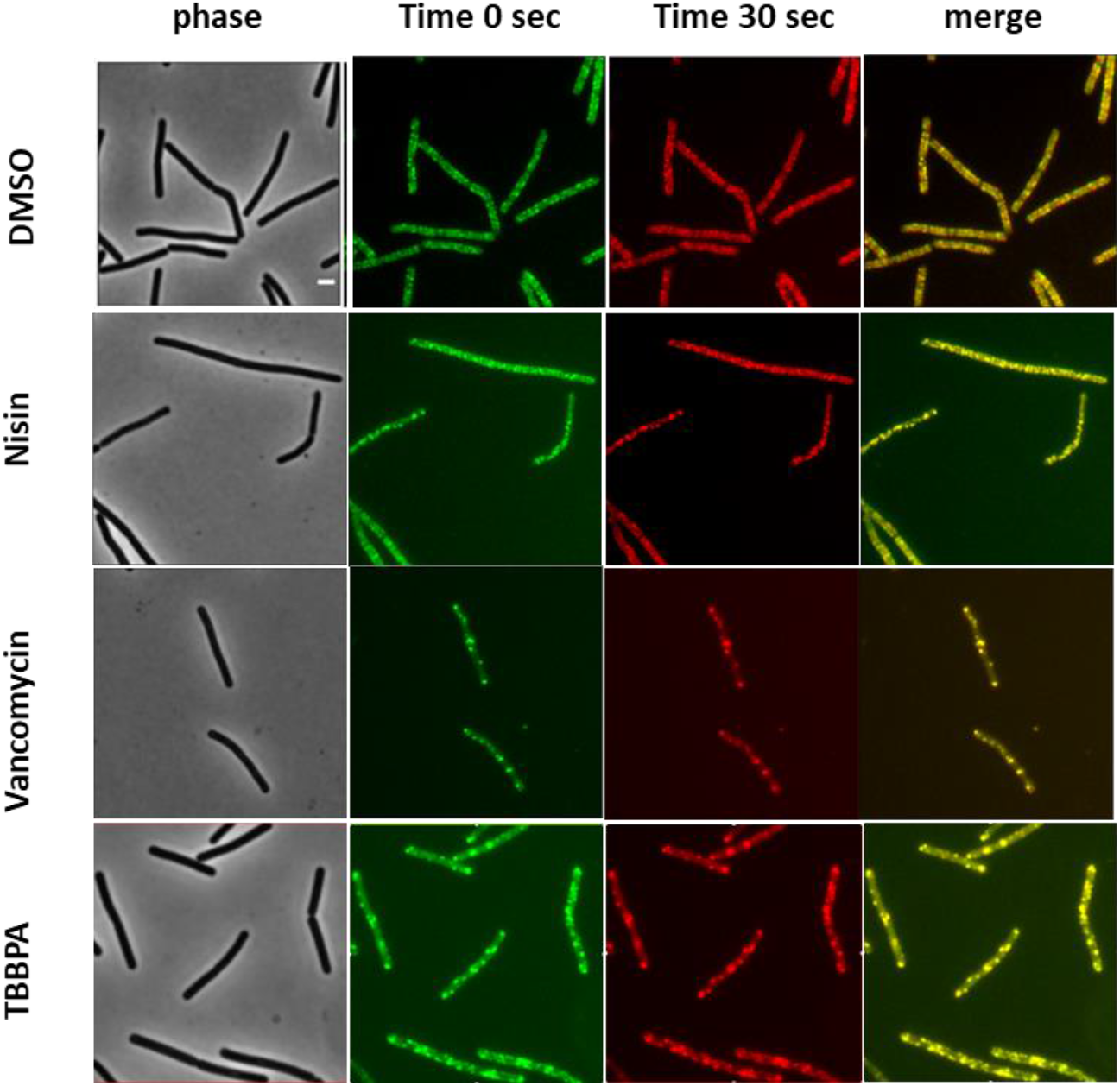
Effect of TBBPA on MreB mobility in *B. subtilis* cells. MreB mobility was evaluated by capturing two separate images of the same *B. subtilis* strain (Pxyl-*gfp-mreB*) cells within a 30-second interval. Individual images were color-coded with red and green, and their overlap indicated stalling of MreB movement (yellow foci). Scale bar 1 μm.

### TBBPA induces membrane depolarization

The fact that TBBPA blocks the movement of MreB might suggest that this compound interferes with peptidoglycan synthesis, leading to the observed cell lysis (Fig. 1B). However, it has been shown that depolarization of the cell membrane disturbs the normal localization of MreB in *B. subtilis* [37]. To examine whether TBBPA affects the membrane potential, we tested the cellular uptake of the membrane potential sensitive probe 3-3′-dipropylthiadicarbocyanine iodide (DiSC3(5)), which is an established reporter for the bacterial membrane potential [38]. However, it appeared that TBBPA directly affects the fluorescence of this probe. So, we tried the fluorescent cationic dye Thioflavin T (ThT). This positively charged dye accumulates in cells with a membrane potential, which increases its fluorescence (Fig. 4A) [39, 40]. The addition of the protonophore carbonyl cyanide m-chlorophenyl hydrazone (CCCP), results in a quick release of ThT from cells (Fig. 4A). Interestingly, the same occurs upon the addition of 8 μg/ml TBBPA (Fig. 4A). To further check that TBBPA dissipates the membrane potential, we made use of a MinD-GFP reporter strain. MinD is a peripheral membrane protein, which localizes at the cell poles and division plane, and is involved in the Min system that prevents aberrant polymerization of the key cell division protein FtsZ close to cell poles in many rod-shaped bacteria [37]. Importantly, its association with the membrane relies on the membrane potential [41]. As illustrated in Fig. 4B, the normal localization of GFP-MinD is completely disturbed when cells are exposed to TBBPA, confirming that the compound causes membrane depolarization.

**Fig. 4.**
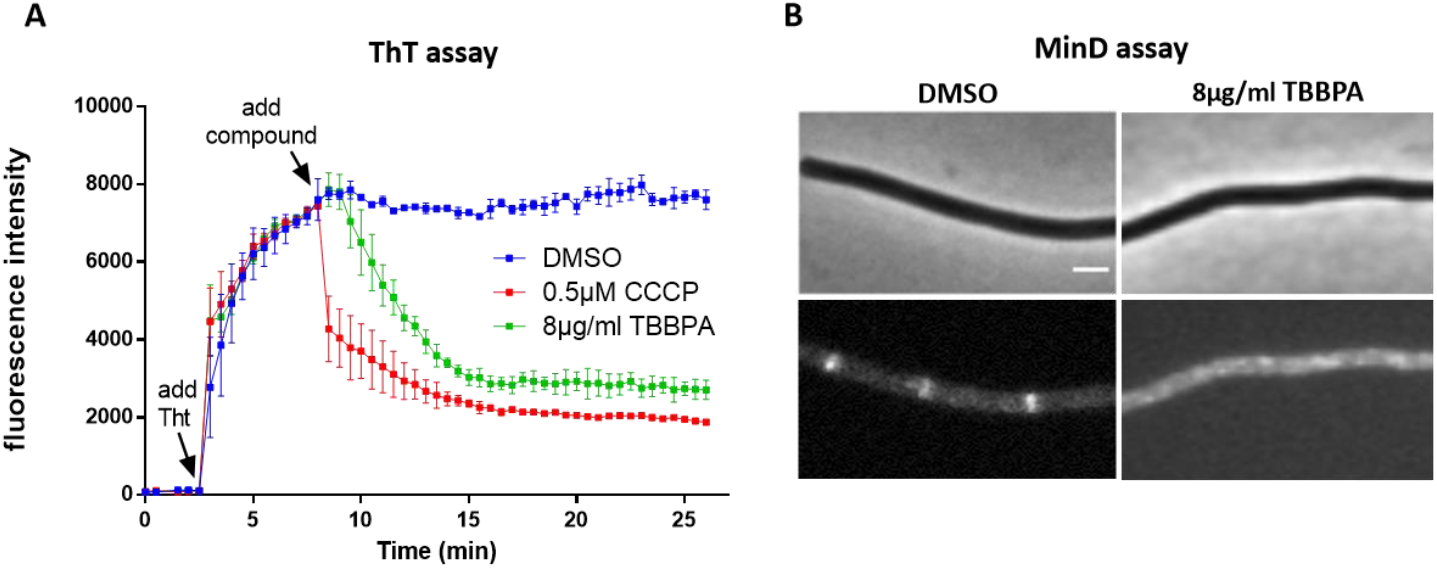
Effects of TBBPA on the membrane potential of *B. subtilis* 168 *cells*. (A) Spectroscopic measurements of membrane potential in *B. subtilis* 168 stained with the fluorescent dye Thioflavin T (ThT); 0.5μM CCCP was used as a positive control. (B) *B. subtilis* strain TB35 (harboring GFP-labeled MinD) was cultured to an OD_600_ of 0.3 and subsequently treated with DMSO (negative control) or 8 μg/ml TBBPA (1x MIC). See also overview microscopy pictures in S1 and S2 Figs. Scale bar 1 μm.

### TBBPA disturbs the cell membrane

Since TBBPA affects the membrane potential, we wondered whether any disturbance of the cell membrane can be observed microscopically. To investigate this, we stained cells with the fluorescent membrane dye FM5-95. As shown in Fig. 5, TBBPA drastically affects the fluorescent membrane stain, inducing prominent fluorescent foci. Such foci may result from the accumulation of extra membrane material or a local increase in membrane fluidity [22, 29, 42]. As a control for intact cells, we also stained the cells with the fluorescence DNA-binding dye 4’,6-diamidino-2-phenylindole (DAPI). Although the DNA was still present in TBBPA treated cells (Fig. 5), the nucleoids appeared strongly condensed.

**Fig. 5.**
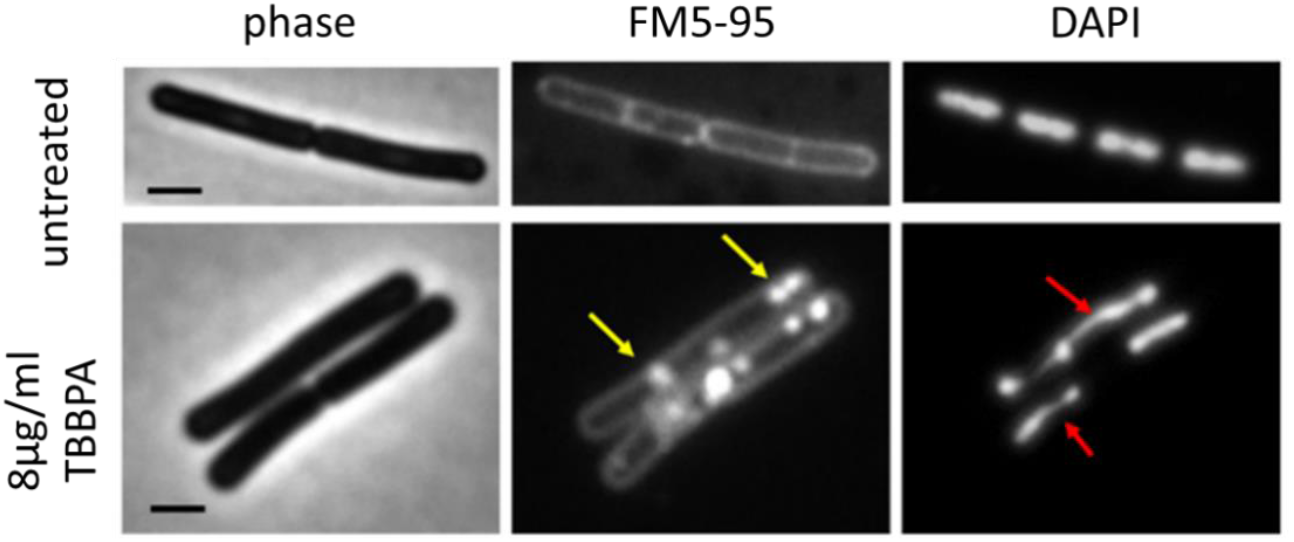
Effects of TBBPA on the membrane and nucleoid of *B. subtilis* 168 cells. *B. subtilis* 168 cells, either untreated or treated with 8 μ g/ml TBBPA, were stained with the membrane-specific dye FM5-95 and the DNA dye DAPI. Yellow arrows indicate TBBPA-induced membrane patches, whereas red arrows point to condensed chromosomes. See also overview microscopy pictures in S3 – S6 Figs. Scale bar 1 μm.

Previous studies have shown that nucleoid condensation can occur with pore-forming compounds such as nisin and tyrocidine [29, 34, 43], presumably because of a loss of turgor pressure [29, 44]. The latter can also cause internal membrane vesical formation due to cell shrinkage, which may explain the fluorescence FM5-95 foci in the cell (Fig. 5). To check whether TBBPA causes membrane perforations, we examined whether cells became permeable for propidium iodide (PI). Propidium iodide shows a fluorescent stain when bound to DNA and RNA, but cannot pass an intact cell membrane, and it is widely used for bacterial viability detection or indicator of membrane integrity [45]. Fig. 6A shows that after 15 min incubation with TBBPA phase dark cells exhibited strong fluorescent staining with propidium iodide, comparable to the addition of 0.5% SDS. This was confirmed with culture measurements in microtiter plates (Fig. 6B), demonstrated that TBBPA severely compromises the cell membrane integrity before cell lysis.

**Fig. 6.**
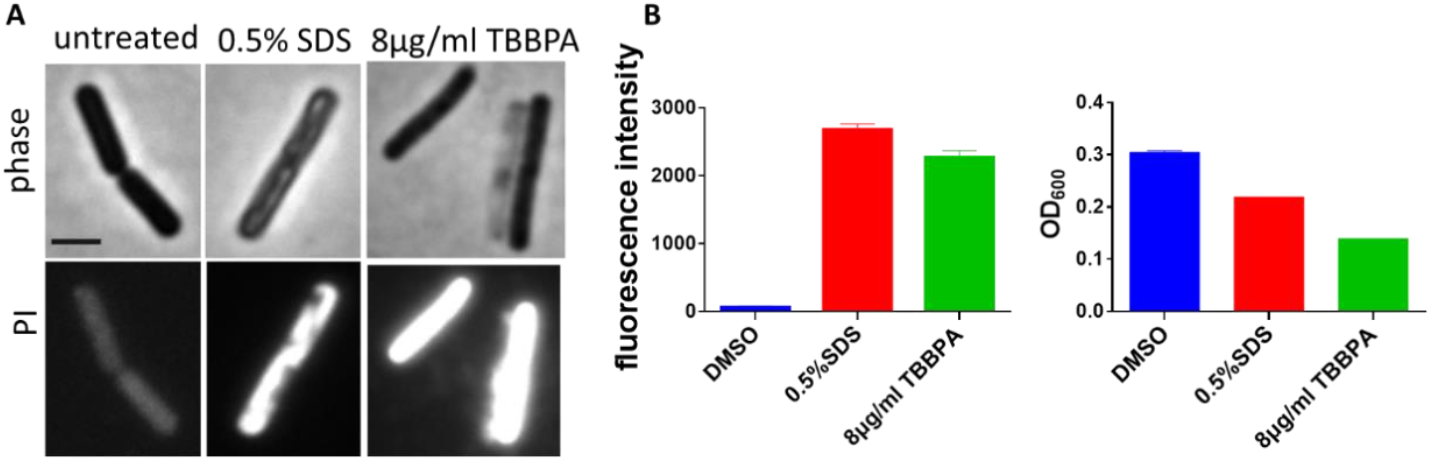
Impact of TBBPA on the integrity of *B. subtilis*. (A) Fluorescence images of *B. subtilis* 168 stained with propidium iodide (PI); 0.5% SDS served as a positive control. (B) Uptake of PI by *B. subtilis* 168 cells treated with the indicated concentrations of SDS or TBBPA; an equivalent amount of DMSO was added as an untreated control. OD_600_ values measured simultaneously with fluorescence. See also overview microscopy pictures in S7 – S9 Figs. Scale bar 1 μm.

### TBBPA affects lipid membranes *in vitro*

TBBPA contains 4 hydrophobic bromine side groups (Fig. 1) and has been shown to partition into artificial lipid membranes resulting in a decrease in membrane fluidity [46]. To examine whether TBBPA can disrupt lipid membranes, we examined its effects on phospholipid bilayers *in vitro* using biomembrane-mimicking giant unilamellar vesicles (GUVs) prepared from *Escherichia coli polar* lipid extracts. The addition of 0.2 % SDS resulted in a shrinkage of GUVs after 5 min, and disappearance within the following minute (Fig. 7). A comparable effect was observed with 80 µg/ml TBBPA (Fig. 7). This finding, together with our other findings and that of others [46], suggests that the bacterial lipid membrane is the target of TBBPA.

**Fig. 7.**
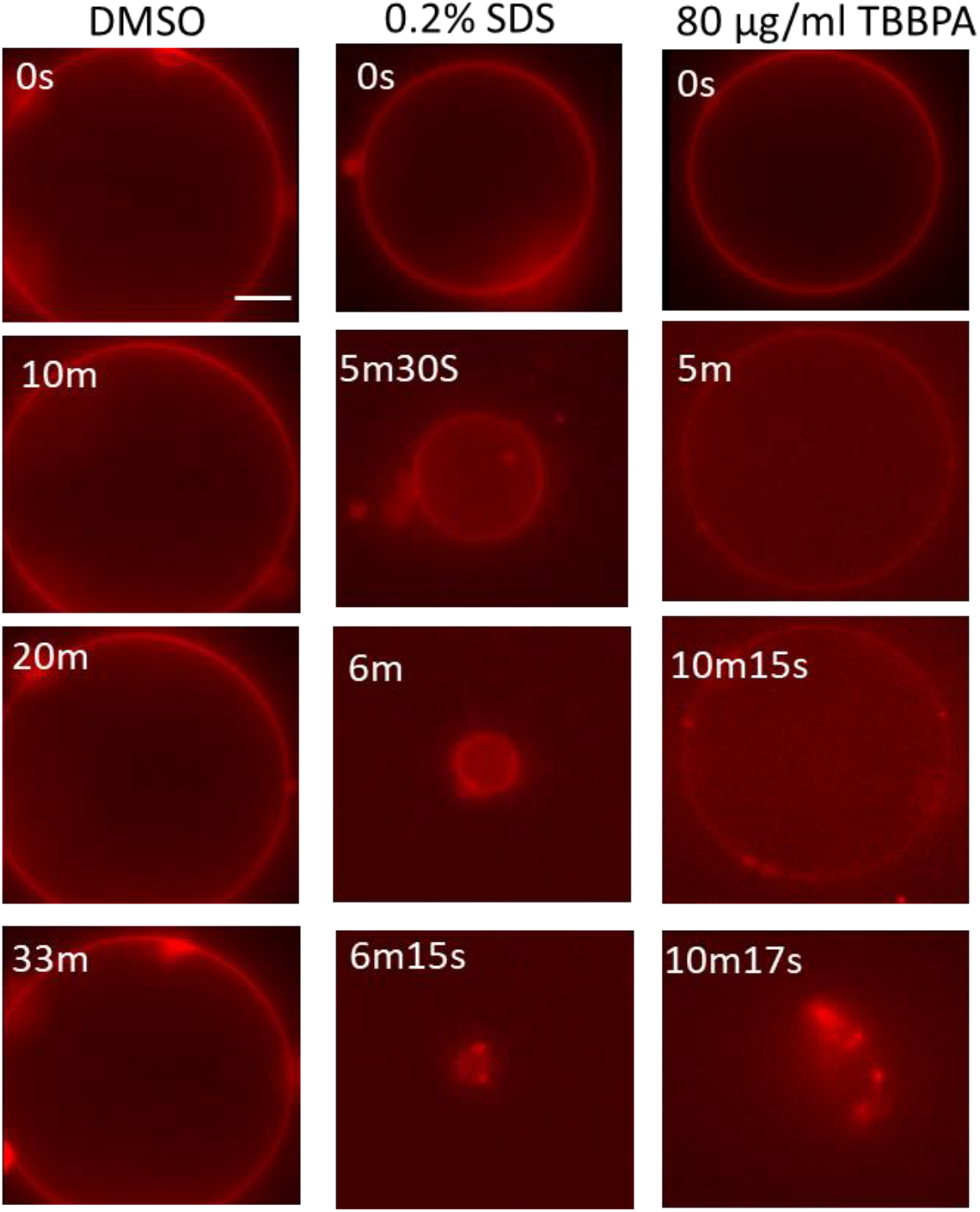
Effect of TBBPA on bacterial lipid bilayers. Giant unilamellar vesicles (GUVs) consisting of *E. coli* polar lipids labeled with 1 μg/ml Nile Red were treated with 80 μg/ml TBBPA or 0.1% DMSO (negative control) and 0.2% SDS (positive control). Observations were made over time using fluorescence microscopy. See also S1-S3 Movies. Scale bar 10 μm.

### Effect of autolysin mutant

The rapid cell lysis following TBBPA exposure is likely due to increased autolysin activity. To test this, we employed a multiple autolysin-deficient mutant strain that lacks 40 cell wall hydrolases [47]. Notably, no obvious cell lysis was detected in such culture, even after 16h of TBBPA treatment (Fig. 8), despite the fact that TBBPA also affected the fluorescence membrane and DAPI stain of this autolysin mutant (not shown). Thus the strong cell lysis was indeed caused by uncontrolled autolysins.

**Fig. 8.**
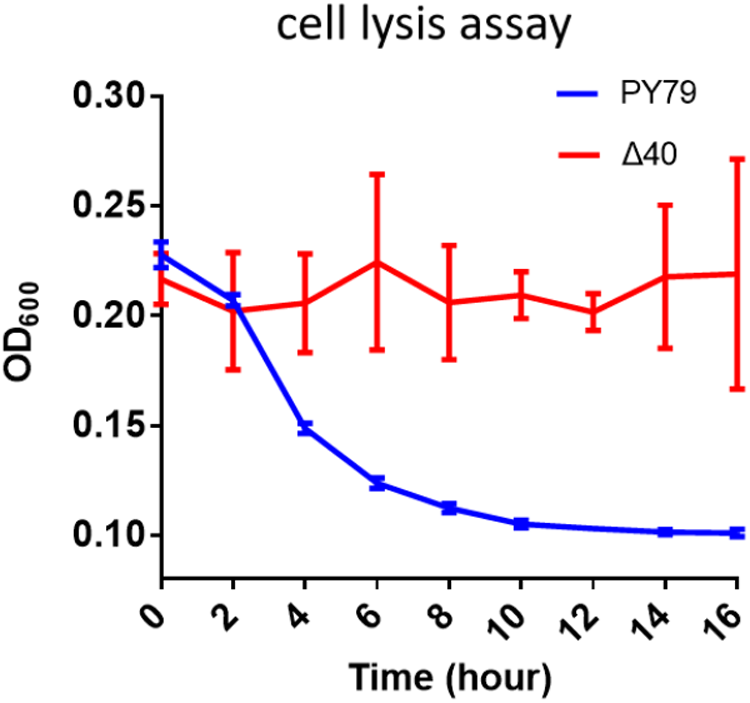
Effects of TBBPA on the cell integrity of *B. subtilis* strains, including the autolysin-deficient mutant Δ40(see S1 Table for detailed information). Changes in OD_600_absorbance of wild-type PY79 and its isogenic mutant after treatment with 16 μg/ml TBBPA.

## DISCUSSION

In this study, we have investigated the antibacterial mechanism of TBBPA against the Gram-positive model bacterium *B. subtilis*. Based on our *in vivo* and *in vitro* experiments and previous studies, we propose that TBBPA directly targets and inserts into the bacterial cell membrane due to its hydrophobic structure [46]. This destabilizes the normal lipid bilayer organization, resulting in the dissipation of the membrane potential, as evidenced by the release of ThT and the abnormal localization of MinD, and eventually formation of large membrane pores, as shown by the entrance of propidium iodine into cells, and the large membrane distortions that can be observed using fluorescence membrane dyes. Cytological profiling revealed that the membrane damage affects a variety of essential processes, including the activity of MreB, indicating that cell wall synthesis is blocked, as well as DNA replication and protein translation. Cells will eventually lyse due to the uncontrolled and/or elevated activity of autolysins.

The induced lysis can be attributed to different mechanisms. Firstly, a lack of peptidoglycan synthesis will distort the normal balance between peptidoglycan polymerization and autolysin-regulated cleavage of peptidoglycan strands, necessary for the insertion of new cell wall material [48, 49]. Secondly, the exposure to TBBPA has been shown to induce the expression of the autolysin genes *yocH* and *cwlO* [2], and thirdly, it has been shown that membrane dissipation itself can lead to uncontrolled activity of autolysins and lysis of *B. subtilis* cells [50]. It has been speculated that this is related to a change in pH gradient in the cell wall, which affects the local activity of autolysins [50]. In any case, the absence of autolysins prevents the lysis normally observed upon treatment with TBBPA.

The selective bactericidal effect of TBBPA against Gram-positive bacteria, with no obvious activity against Gram-negative counterparts, is most likely attributable to the outer membrane, which serves as a robust physical barrier. The densely packed lipopolysaccharides in the outer membrane form a hydrophilic surface hydration layer that encases the cell, restricting the accessibility of hydrophobic compounds such as TBBPA [51].

TBBPA holds considerable potential for treating drug-resistant Gram-positive bacterial infections including MRSA, primarily due to its low propensity to induce drug resistance. This advantage is plausibly linked to its membrane-targeting mechanism, which is difficult to alter, unlike conventional antibiotics targeting specific enzymes/receptors [52-54]. Nevertheless, TBBPA’s clinical translation is constrained by unresolved environmental toxicity and human safety concerns, likely related to its membrane disruptive activity, which demand rigorous evaluation. Future research should focus on structural optimization of TBBPA to enhance antibacterial activity while mitigating toxicity. Previous structure-activity relationship studies have indicated that TBBPA’s antimicrobial efficacy is closely associated with its halogen substitutions and phenolic hydroxyl groups [2], providing a rational basis for chemical modifications (e.g., adjusting bromine substituents, introducing hydrophilic moieties) or conjugation with Gram-positive targeting ligands to improve selective activity. Ideally, such modifications lead to a preference for targeting and interfering with bacterial membranes instead of eukaryotic membranes. Additionally, TBBPA-based nanodelivery systems may enhance solubility, extend circulation, and enable site-specific delivery to infection foci. With advancements in medicinal chemistry, optimized TBBPA derivatives are expected to overcome safety barriers, unlocking their potential as novel therapeutics against drug-resistant Gram-positive infections.

## ACKNOWLEDGEMENTS

We are grateful to Professor Ethan Garner from Harvard University for generously providing a series of *Bacillus subtilis* strains deficient in various hydrolases. We thank Dr. Haibo Liu for his helpful advice, insightful discussions, and critical reading of the manuscript. This work was financially supported by the China Scholarship Council (CSC) (No.202208420147).

## REFERENCES

1. Murray, C.J.L., et al., Global burden of bacterial antimicrobial resistance in 2019: a systematic analysis. The Lancet, 2022. 399(10325): p. 629–655.

2. Ji, F., et al., Tetrabromobisphenol A (TBBPA) exhibits specific antimicrobial activity against Gram-positive bacteria without detectable resistance. Chem Commun (Camb), 2017. 53(25): p. 3512–3515.

3. Segev, O., A. Kushmaro, and A. Brenner, Environmental impact of flame retardants (persistence and biodegradability). Int J Environ Res Public Health, 2009. 6(2): p. 478–91.

4. National Center for Biotechnology Information. PubChem Compound Summary for CID 6618, Tetrabromobisphenol A. 2025 January 15, 2025; Available from: https://pubchem.ncbi.nlm.nih.gov/compound/Tetrabromobisphenol-A.

5. Miao, B., et al., A Review on Tetrabromobisphenol A: Human Biomonitoring, Toxicity, Detection and Treatment in the Environment. Molecules, 2023. 28(6).

6. Hu, J., et al., Assessing the toxicity of TBBPA and HBCD by zebrafish embryo toxicity assay and biomarker analysis. Environ Toxicol, 2009. 24(4): p. 334–42.

7. Kitamura, S., et al., Thyroid hormonal activity of the flame retardants tetrabromobisphenol A and tetrachlorobisphenol A. Biochem Biophys Res Commun, 2002. 293(1): p. 554–9.

8. Levy-Bimbot, M., et al., Tetrabromobisphenol-A disrupts thyroid hormone receptor alpha function in vitro: use of fluorescence polarization to assay corepressor and coactivator peptide binding. Chemosphere, 2012. 87(7): p. 782–8.

9. Hurd, T. and M.M. Whalen, Tetrabromobisphenol A decreases cell-surface proteins involved in human natural killer (NK) cell-dependent target cell lysis. J Immunotoxicol, 2011. 8(3): p. 219–27.

10. Strack, S., et al., Cytotoxicity of TBBPA and effects on proliferation, cell cycle and MAPK pathways in mammalian cells. Chemosphere, 2007. 67(9): p. S405–11.

11. Tada, Y., et al., Flame retardant tetrabromobisphenol A induced hepatic changes in ICR male mice. Environmental Toxicology and Pharmacology, 2007. 23(2): p. 174–178.

12. International Agency for Research on Cancer. IARC Monographs on the Identification of Carcinogenic Hazards to Humans. 2018; Available from: https://monographs.iarc.who.int/list-of-classifications.

13. Grasselli, E., et al., Thyromimetic actions of tetrabromobisphenol A (TBBPA) in steatotic FaO rat hepatoma cells. Chemosphere, 2014. 112: p. 511–8.

14. Oral, D., et al., Toxic Effects of Tetrabromobisphenol A: Focus on Endocrine Disruption. J Environ Pathol Toxicol Oncol, 2021. 40(3): p. 1–23.

15. European Chemicals Bureau. European Union Risk Assessment Report 2,2’,6,6’-tetrabromo-4,4’-isopropylidenediphenol (tetrabromobisphenol-A or TBBP-A) Part II – human health. 2006; Available from: https://echa.europa.eu/documents/10162/32b000fe-b4fe-4828-b3d3-93c24c1cdd51.

16. Zhou, H., N. Yin, and F. Faiola, Tetrabromobisphenol A (TBBPA): A controversial environmental pollutant. J Environ Sci (China), 2020. 97: p. 54–66.

17. Wecke, J., M. Perego, and W. Fischer, D-alanine deprivation of Bacillus subtilis teichoic acids is without effect on cell growth and morphology but affects the autolytic activity. Microb Drug Resist, 1996. 2(1): p. 123–9.

18. Pucci, M.J., L.F. Discotto, and T.J. Dougherty, Cloning and identification of the Escherichia coli murB DNA sequence, which encodes UDP-N-acetylenolpyruvoylglucosamine reductase. J Bacteriol, 1992. 174(5): p. 1690–3.

19. Popham, D.L. and P. Setlow, Cloning, nucleotide sequence, and mutagenesis of the Bacillus subtilis ponA operon, which codes for penicillin-binding protein (PBP) 1 and a PBP-related factor. J Bacteriol, 1995. 177(2): p. 326–35.

20. Murray, T., D.L. Popham, and P. Setlow, Identification and characterization of pbpA encoding Bacillus subtilis penicillin-binding protein 2A. J Bacteriol, 1997. 179(9): p. 3021–9.

21. Nonejuie, P., et al., Bacterial cytological profiling rapidly identifies the cellular pathways targeted by antibacterial molecules. Proc Natl Acad Sci U S A, 2013. 110(40): p. 16169–74.

22. Muller, A., et al., Daptomycin inhibits cell envelope synthesis by interfering with fluid membrane microdomains. Proc Natl Acad Sci U S A, 2016. 113(45): p. E7077–E7086.

23. Hu, Y., et al., A new approach for the discovery of antibiotics by targeting non-multiplying bacteria: a novel topical antibiotic for staphylococcal infections. PloS one, 2010. 5(7): p. e11818.

24. Weinberger, A., et al., Gel-Assisted Formation of Giant Unilamellar Vesicles. Biophysical Journal, 2013. 105(1): p. 154–164.

25. Zhu, Y., et al., Gene clusters located on two large plasmids determine spore crystal association (SCA) in Bacillus thuringiensis subsp. finitimus strain YBT-020. PloS one, 2011. 6(11): p. e27164.

26. Sharpe, M.E., et al., Bacillus subtilis cell cycle as studied by fluorescence microscopy: constancy of cell length at initiation of DNA replication and evidence for active nucleoid partitioning. J Bacteriol, 1998. 180(3): p. 547–55.

27. Su’etsugu, M. and J. Errington, The replicase sliding clamp dynamically accumulates behind progressing replication forks in Bacillus subtilis cells. Mol Cell, 2011. 41(6): p. 720–32.

28. Lewis, P.J., S.D. Thaker, and J. Errington, Compartmentalization of transcription and translation in Bacillus subtilis. Embo j, 2000. 19(4): p. 710–8.

29. Wenzel, M., et al., The Multifaceted Antibacterial Mechanisms of the Pioneering Peptide Antibiotics Tyrocidine and Gramicidin S. MBio, 2018. 9(5).

30. Eraso, J.M. and W. Margolin, Bacterial cell wall: thinking globally, actin locally. Curr Biol, 2011. 21(16): p. R628–30.

31. Domínguez-Escobar, J., et al., Processive movement of MreB-associated cell wall biosynthetic complexes in bacteria. Science, 2011. 333(6039): p. 225–8.

32. Garner, E.C., et al., Coupled, circumferential motions of the cell wall synthesis machinery and MreB filaments in B. subtilis. Science, 2011. 333(6039): p. 222–5.

33. van Teeffelen, S., et al., The bacterial actin MreB rotates, and rotation depends on cell-wall assembly. Proc Natl Acad Sci U S A, 2011. 108(38): p. 15822–7.

34. Schäfer, A.B., et al., Dissecting antibiotic effects on the cell envelope using bacterial cytological profiling: a phenotypic analysis starter kit. Microbiol Spectr, 2024: p. e0327523.

35. Wiedemann, I., R. Benz, and H.G. Sahl, Lipid II-Mediated Pore Formation by the Peptide Antibiotic Nisin: a Black Lipid Membrane Study. J Bacteriol, 2004. 186(10): p. 3259–3261.

36. Reynolds, P.E., Structure, biochemistry and mechanism of action of glycopeptide antibiotics. Eur J Clin Microbiol Infect Dis, 1989. 8(11): p. 943–50.

37. Müller, A., et al., Daptomycin inhibits cell envelope synthesis by interfering with fluid membrane microdomains. Proc Natl Acad Sci U S A, 2016. 113(45): p. E7077–e7086.

38. Skates, E., et al., Thioflavin T indicates mitochondrial membrane potential in mammalian cells. Biophys Rep (N Y), 2023. 3(4): p. 100134.

39. Prindle, A., et al., Ion channels enable electrical communication in bacterial communities. Nature, 2015. 527(7576): p. 59–63.

40. Stratford, J.P., et al., Electrically induced bacterial membrane-potential dynamics correspond to cellular proliferation capacity. Proc Natl Acad Sci U S A, 2019. 116(19): p. 9552–9557.

41. Strahl, H. and L.W. Hamoen, Membrane potential is important for bacterial cell division. Proc Natl Acad Sci U S A, 2010. 107(27): p. 12281–6.

42. Strahl, H., F. Bürmann, and L.W. Hamoen, The actin homologue MreB organizes the bacterial cell membrane. Nat Commun, 2014. 5: p. 3442.

43. Jensen, C., et al., Nisin Damages the Septal Membrane and Triggers DNA Condensation in Methicillin-Resistant Staphylococcus aureus. Front Microbiol, 2020. 11: p. 1007.

44. Cagliero, C. and D.J. Jin, Dissociation and re-association of RNA polymerase with DNA during osmotic stress response in Escherichia coli. Nucleic Acids Res, 2013. 41(1): p. 315–26.

45. Rosenberg, M., N.F. Azevedo, and A. Ivask, Propidium iodide staining underestimates viability of adherent bacterial cells. Scientific Reports, 2019. 9(1): p. 6483.

46. Ogunbayo, O.A., K.T. Jensen, and F. Michelangeli, The interaction of the brominated flame retardant: tetrabromobisphenol A with phospholipid membranes. Biochim Biophys Acta, 2007. 1768(6): p. 1559–66.

47. Wilson, S.A., et al., An exhaustive multiple knockout approach to understanding cell wall hydrolase function in Bacillus subtilis. mBio, 2023. 14(5): p. e0176023.

48. Vollmer, W., et al., Bacterial peptidoglycan (murein) hydrolases. FEMS Microbiol Rev, 2008. 32(2): p. 259–86.

49. Uehara, T. and T.G. Bernhardt, More than just lysins: peptidoglycan hydrolases tailor the cell wall. Curr Opin Microbiol, 2011. 14(6): p. 698–703.

50. Jolliffe, L.K., R.J. Doyle, and U.N. Streips, The energized membrane and cellular autolysis in Bacillus subtilis. Cell, 1981. 25(3): p. 753–63.

51. Konovalova, A., D.E. Kahne, and T.J. Silhavy, Outer Membrane Biogenesis. Annu Rev Microbiol, 2017. 71: p. 539–556.

52. Lety, M.A., et al., A single point mutation in the embB gene is responsible for resistance to ethambutol in Mycobacterium smegmatis. Antimicrob Agents Chemother, 1997. 41(12): p. 2629–33.

53. Musser, J.M., et al., Reduced In Vitro Susceptibility of Streptococcus pyogenes to β-Lactam Antibiotics Associated with Mutations in the pbp2x Gene Is Geographically Widespread. J Clin Microbiol, 2020. 58(4).

54. Sánchez-Maroto, L., et al., Idiosyncratic evolvability among single-point ribosomal mutants towards multi-aminoglycoside resistance. PLoS Genet, 2025. 21(8): p. e1011832.

